# Exogenously applied ABA induced changes in physio-biochemical attributes in selected rice varieties under osmotic stress

**DOI:** 10.1101/2020.05.27.116129

**Authors:** Goutam Kumar Dash, Chaitanya Kumar Geda, Soumya Kumar Sahoo, Mirza Jaynul Baig, Padmini Swain

## Abstract

Drought is the most catastrophic abiotic stress that affect plant growth and development. Plant initiates alterations at physiological, biochemical and molecular levels to combat deleterious effects of drought in which abscisic acid, play a pivotal role. The present investigation was conducted to investigate how abscisic acid modulates different biochemical traits under osmotic stress. Twenty-one days old seedlings were exposed to osmotic stress, exogenously applied abscisic acid and the combination of both osmotic stress and abscisic acid to study the influence of abscisic acid on biochemical traits under moisture stress. The results reveals that abscisic acid has no influence on leaf water potential, however, it has active role in increasing osmolytes like proline. Δ^1^-pyrroline-5-carboxylate reductase and proline dehydrogenase remained unresponsiveness to ABA that indicates ABA-independent regulation of these two enzymes but the activity of Δ^1^-pyrroline-5-carboxylate synthesase and Ornithine-δ-aminotransferase was enhanced (3 fold) in response to abscisic acid. Antioxidative enzymes increased (catalase-2.5 fold; peroxidase-3 fold and SOD-6 fold) in response to abscisic acid and osmotic stress, gave an insight of the regulation of these traits under the influence of abscisic acid. Aldehyde oxidase remained unresponsive to abscisic acid but showed enhanced expression when both abscisic acid and osmotic stress was applied suggest its ABA-independent regulation. The information provided here has significance in understanding the regulation of different catabolite that play important role in drought tolerance and can be implemented for the development of drought-tolerant varieties.

## Introduction

Drought is the most significant environmental stress in agriculture that affect plant growth and productivity (Xoconostle-Cazares et al. 2010). It impairs physiological and biochemical processes and causes severe damage to the plant. To nullify the effect of drought, plants develop certain morphological (Farooq et al. 2010; Dash et al. 2017), physiological (Sibounheuang et al. 2006), biochemical and molecular mechanisms by altering its cellular metabolism and defense mechanism that includes closing of stomata to reduce water loss, accumulation of osmo-regulators to retain water inside the cell and production of reactive oxygen species (ROS) scavenging enzymes to neutralize superoxide radicals etc. The phytohormone abscisic acid plays an important role in drought tolerance mechanism which is accumulated mainly in response to drought stress. Increase in endogenous abscisic acid (ABA) level with the commencement of water stress and gradually degradation upon release of stress has been well documented (Zhang et al. 2006). ABA is thought to be an important signalling factor involved in drought tolerance. Plants having high endogenous ABA level are considered to reduce water loss through transpiration and exhibit efficient drought tolerance mechanism (Kholová et al. 2010). Evidence of growth promotion and an increase in leaf water content by foliar spraying of ABA under drought condition has been documented (Sansberro et al. 2004). ABA enhances leaf relative water content through a simultaneous increase in water uptake by aquaporins at root level and transpiration check at leaf level (Lian et al. 2006). Besides activating aquaporins, ABA increases accumulation of osmo-regulators that help in lowering water potential of the cell thus generating water potential gradient that drives uptake of water from the soil even when stomata are closed and production of anti-oxidative enzymes that protect the essential proteins and photosystems against desiccation injury (Dash and Swain 2015). Proline and soluble sugars are common osmo-regulator that are accumulated in response to drought (Jiang et al. 2012). (Savoure et al. (1997) reported an increase in P5CS gene expression under water stress in response to 50 μM ABA but P5CR gene expression was scarcely detectable in response to ABA. Proline dehydrogenase (PDH) that is responsible for degradation of proline has been reported to decrease under water stress condition (Sharma and Verslues 2010). Proline is also synthesized in an alternate pathway using ornithine as precursors through the enzyme ornithine-δ-aminotransferases (OAT). Expression of Ornithine-δ-aminotransferase (OAT) in an ABA-dependent and ABA-independent manner has also been reported (You et al. 2012). Similarly, sugar accumulation in response to water stress contributes in maintaining osmotic adjustment. Increase in total sugar and reducing sugars under water stress and in response to ABA has been observed with increased α- and β- amylase activity (Sarfaraz-Ardakani et al. 2014). During water stress, many authors have reported increased synthesis of anti-oxidative enzymes. (Jiang and Zhang 2001) reported increased generation of O_2_ and induction of antioxidant enzyme gene expression in response to exogenously applied. But reports regarding the effect of ABA on its own biosynthesis enzymes like aldehyde oxidase (AO) is limited. In the present study, attempts are made to characterise some identified drought tolerant and susceptible genotypes on the basis of ABA modulated physiological and biochemical traits during osmotic stress.

## Materials and Methods

### Plant material

Four previously identified promising rice genotypes (*Oryza sativa* L.) (AC 43037, AC 43025, AC 43012 and Lalaijung) seems to be tolerant to drought stress at vegetative stage along with one tolerant (CR 143-2-2) & susceptible check (IR 64) were taken in the experimentation to further detailed study. Seeds were sterilized with 10% sodium hypochlorite solution and then germinated in petri dishes on moist filter paper. Sprouting seeds were placed in the holes of polystyrene plates supported by nylon mess after two days of germination. The polystyrene plates were placed on the surface of plastic trays containing 10 lt. of full-strength nutrient medium as described by Yoshida et al. 1976. Three sets of trays were arranged to maintain three replications (12 no. of trays) per treatment and the seedling of all varieties were placed in each tray in separate rows. Plants were allowed to grow for 21 days in the nutrient solution and pH of the nutrient solution was maintained at 5.7 every day. Nutrient solution was renewed in every 5 days interval. After 21 days, the seedlings were exposed to four types of treatments for 6 days to study the effect of ABA on different biochemical traits in the presence as well as in the absence of osmotic stress. The treatments were as follows:

1. medium containing only nutrient solution i.e. control (C)
2. medium containing nutrient solution along with 2% D-mannitol (M)
3. medium containing nutrient solution along with 10 μM ABA (ABA)
4. medium containing nutrient solution, 2% D-mannitol and 10 μM ABA (M+ABA)

For each treatment, five replications were made. Osmotic potential of 2% D-mannitol was estimated to be 0.59 MPa using water potential system, WESCOR, USA. Leaf sampling was done at sixth day of treatments applied. Fully expanded second leaf was collected in a polyethylene bag at the midday (1.00-2.00 pm) for estimation of physiological and biochemical traits and were stored at −80 °C.

### Determination of Leaf water potential (LWP)

Leaf water potential was measured following the method of H.D. Barrs and P.E. Weatherley 1962 and Turner 1982 with the help of water potential system, Psypro, Wescor (USA).

### Estimation of Proline content

Total proline content was estimated following (Bates et al. 1973). Fresh leaf sample of 0.5 g was homogenized in 10 ml of 3% aqueous sulfosalicylic acid and centrifuged at 10,009 g for 10 min at 4 °C. Two millilitres of the supernatant was mixed with 2 ml of glacial acetic acid and 2 ml of acid ninhydrin (1.25 g ninhydrin in 30 ml glacial acetic acid and 20 ml 6 M phosphoric acid) and kept in boiling water bath for 1 hr. The reaction was stopped by placing the tubes in an ice bath. After cooling to room temperature, 4 ml toluene was added to the reaction mixture and vortex shaken. The toluene layer was separated and the red colour intensity was measured at 520 nm.

### Enzyme assay for proline metabolism

Enzyme extraction from the frozen leaf samples was carried out according to Lutts et al. 1999. One gram frozen leaf sample was homogenized in 100 mM potassium phosphate buffer containing 1 mM EDTA, 10 mM β-mercaptoethanol, 1% polyvinylpolypyrrolidone, 5 mM MgCl_2_ and 0.6 M KCl maintained at pH 7.4. The homogenate was centrifuged at 13,000 g for 15 min at 4 °C.

The activity of Δ^1^-pyrroline-5-carboxylate synthase (P5CS; EC 2.7.2.11) was determined according to the method of Stines et al. (1999). Reaction was started by adding 0.5 ml of enzyme extract to a mixture of 75 mM L-glutamate, 20 mM MgCl_2_, 100 mM Tris–HCl, 5 mM ATP and 0.4 mM NADPH. After incubation for 20 min at 37 °C, absorbance was recorded at 340 nm using UV-Visible spectrophotometer (Thermo Fisher Scientific, Finland). Enzyme activity was expressed as U mg protein^−1^ considering 1 U is equivalent to an increase of 0.001 A_340_ per min.

Δ^1^-pyrroline-5-carboxylate reductase (P5CR; EC 1.5.1.2) activity was measured following the method of Madan et al. (1995). To a mixture of 0.06 mM NADH, 0.15 mM Δ^1^-pyrroline-5-carboxylic acid, 120 mM potassium phosphate buffer and 2 mM dithiothreitol, 0.5 ml of enzyme extract was added and the decrease in the absorbance was recorded at 340 nm. P5CR activity was expressed as U mg protein^−1^ (one unit defined as a decrease in 0.01 A_340_ per min).

Ornithine-δ-aminotransferase (OAT; EC 2.6.1.68) was assayed by the method of Vogel and Kopac (1960). Enzyme extract and 100 mM potassium phosphate buffer (pH 8.0) was mixed in the proportion 0.2:0.8 ml. Buffer contained 50 mM L-ornithine, 20 mM α-ketoglutarate and 1 mM pyridoxal-5-phosphate. The reaction was stopped by adding 0.5 ml 10% trichloro acetic acid and 0.5% O-aminobenzaldehyde in ethanol to the reaction mixture after incubation for 30 min at 37 °C. After 1 hour, the reaction mixture was centrifuged at 12 0009 g for 10 min at 4 °C. Increase in absorbance was recorded at 440 nm. Enzyme activity was expressed as U mg protein^−1^ (one unit defined as an increase in 0.01 A_440)._

For proline dehydrogenase (PDH; EC 1.5.99.8) estimation, the enzyme extract was dissolved in 0.15 M dm^−3^ Na_2_CO_3_-HCl buffer (pH 10.3) with 13 mM dm^−3^ L-proline and 1.5 mM dm^−3^ NAD^+^ ((Lutts et al. 1999) and absorbance was measured at 340 nm. PDH was expressed as U mg protein^−1^ (one unit is defined as an increase in 0.01 A_340_ per min).

### Anti-oxidative enzyme Assay

Leaf material (1 g) was homogenized in 1.5 ml of potassium phosphate buffer (pH 7.8) containing 1 mM EDTA, 1 mM ascorbate, 10% w/v sorbitol and 0.1% Triton X. The homogenate was centrifuged at 15,000g for 20 mins at 4 °C and the supernatant was used for catalase, peroxidase, and superoxide dismutase enzyme analysis. Protein concentration was detected according to the method of (Lowry et al. 1951) using the bovine serum as a standard.

Catalase (EC 1.11.1.6) activity was estimated by the method followed by Pereira et al. 2002. To a mixture of 50 mM potassium phosphate buffer (pH 7.0) and 0.036% of H_2_O_2,_ 0.1 ml of enzyme extract was added and the decrease in absorbance was recorded at 240 nm and the rate of enzyme activity was expressed as U mg protein^−1^.

Peroxidase (EC 1.11.1.x) activity was estimated following the method Pütter (1974). To a mixture of 0.1 M potassium phosphate buffer (pH 7.0) and 0.018 M guaiacol, 0.1 ml of enzyme solution was added. The increase in absorbance was recorded at 436 nm and the rate of enzyme activity was expressed as U mg protein^−1^.

The activity of superoxide dismutase (SOD) (EC 1.15.1.1) was estimated by the method Giannopolitis and Ries (1977). To a mixture of 0.05 M phosphate buffer, 13 mM methionine, 75 μM NBT and 100 nM EDTA, 0.1 ml of enzyme extract was added and placed 30 cm below a light bank consisting of two 30–watt-fluorescent bulbs (reference tube must be placed in the dark). The reaction was started by switching on the light and allowed to run for 20 min and then terminated by switching off the light followed by covering the tubes with a black cloth. Each tube was wrapped with aluminum foil. Absorbance was read against the blank (reference tube) at 560 nm. The volume of enzyme extract producing 50% inhibition of the reaction is defined to have one unit of SOD activity.

### Assay of Aldehyde oxidase (AO) activity

Aldehyde oxidase (AO; EC 1.2.3.1)) activity was determined following the method of Zdunek and Lips (2001). Plant tissue was homogenized immediately after harvesting with acid-washed sand and ice-cold extraction medium containing 250 mM TRIS-HCl (pH 8.5), 1 mM EDTA, 10 mM reduced glutathione (GSH), and 2 mM dithiothreitol (DTT). A ratio of 1 g tissue to 3 ml buffer (1:3 w/v) was used to homogenize plant material and was centrifuged at 27,000g and 4 °C for 15 min. The resulting supernatant was subjected to native polyacrylamide gel electrophoresis (PAGE) with 7.5% polyacrylamide gel in a Laemmli buffer system (Laemmli 1970) in the absence of sodium dodecyl sulphate at 4 °C. Each lane in the gel was loaded with 400 μg leaf proteins. After electrophoresis, AO activity staining was developed at room temperature in a mixture containing 0.1 M TRIS-HCl, pH 7.5, 0.1 mM phenazine methosulphate, 1 mM MTT (3[4,5-dimethylthiazol-2-yl]-2,5-diphenyltetrazolium-bromide) and 1 mM indole-3-aldehyde (substrate). The gel was incubated in dark overnight for the development of bands. The activity of AO was estimated based on MTT reduction, which resulted in the development of specific formazon bands which were directly proportional to enzyme activity (Rothe 1974) and quantified using the NIH Image J 1.6 computer software.

## Statistical Analysis

Analysis of Variance was analysed using Cropstat for windows 7.2.2007.3 (IRRI, Philippines). The least significant difference test was used to distinguish among individual mean values where applicable with a confidence level of *p* < 0.01 and *p* < 0.001.

## Results

### Effect of osmotic stress and ABA on Leaf water potential

Leaf water potential substantially decreased till six days of treatment under osmotic stress and osmotic stress along with ABA (p<0.05) compared to control (C). Exogenous application of ABA did not affect LWP and there was no significant variation in LWP between the control plants and the plants treated with only ABA. LWP also did not seem to be affected under osmotic stress by ABA treatment which was evident from the insignificant difference in LWP between M (−2.11 MPa) and M+ABA (−2.15 MPa) treatment. However, a significant difference in LWP (p<0.05) was observed between tolerant and susceptible genotypes. Decrease in LWP in susceptible genotypes (Lalaijung and IR 64) was more than two fold to that of tolerant genotypes (AC 43025, AC 43012, AC 43037 and CR 143-2-2) both under M and M+ABA treatments (Table 1; Fig 1). LWP did not vary between M and M+ABA treatments, despite the presence of ABA in M+ABA treatment in all the susceptible genotypes.

**Table-1.** Mean of all biochemical traits of six rice genotypes measured under four treatments.

| Traits | Treatments | 6 DAT | Traits | Treatments | 6 DAT |
| --- | --- | --- | --- | --- | --- |
| LWP (-MPa) | C | -1.05 ± 0.04 | PDH (U mg protein<sup>-1</sup>) | C | 4.42 ± 0.15 |
|  | M | -2.12 ± 0.36 |  | M | 1.08 ± 0.10 |
|  | ABA | -1.22 ± 0.05 |  | ABA | 3.52 ± 0.11 |
|  | M+ABA | -2.14 ± 0.34 |  | M+ABA | 1.08 ± 0.21 |
| Proline (µM g DW<sup>-1</sup>) | C | 6.6 ± 0.60 | CAT (U mg protein<sup>-1</sup>) | C | 0.28 ± 0.01 |
|  | M | 148.6 ± 30.25 |  | M | 0.69 ± 0.06 |
|  | ABA | 63.0 ± 7.21 |  | ABA | 0.73 ± 0.03 |
|  | M+ABA | 186.0 ± 19.24 |  | M+ABA | 0.77 ± 0.03 |
| P5CS (U mg protein<sup>-1</sup>) | C | 0.05 ± 0.00 | POX (U mg protein<sup>-1</sup>) | C | 2.43 ± 0.11 |
|  | M | 1.09 ± 0.23 |  | M | 8.08 ± 0.95 |
|  | ABA | 1.04 ± 0.16 |  | ABA | 8.87 ± 0.97 |
|  | M+ABA | 1.35 ± 0.17 |  | M+ABA | 9.33 ± 1.18 |
| P5CR (U mg protein<sup>-1</sup>) | C | 0.03 ± 0.00 | SOD (U mg protein<sup>-1</sup>) | C | 0.46 ± 0.02 |
|  | M | 0.27 ± 0.04 |  | M | 2.53 ± 0.40 |
|  | ABA | 0.04 ± 0.00 |  | ABA | 2.40 ± 0.37 |
|  | M+ABA | 0.27 ± 0.04 |  | M+ABA | 2.77 ± 0.40 |
| OAT (U mg protein<sup>-1</sup>) | C | 0.31 ± 0.01 |  |  |  |
|  | M | 1.02 ± 0.14 |  |  |  |
|  | ABA | 1.29 ± 0.14 |  |  |  |
|  | M+ABA | 1.58 ± 0.14 |  |  |  |

C-Medium containing only nutrient solution i.e. control; M-Medium containing nutrient solution along with 2% D- mannitol to study the effect of osmotic stress on biochemical traits; ABA-Medium containing nutrient solution along with 60 µM ABA to study the effect of ABA on biochemical traits; M+ABA-Medium containing nutrient solution, 2% D-mannitol and 60 µM ABA to study how ABA modulates biochemical traits under osmotic stress; P5CS- Δ1-pyrroline-5-carboxylate synthase; P5CR- Δ1-pyrroline-5-carboxylate reductase; OAT-Ornithine-δ-aminotransferase; PDH-Proline dehydrogenase. CAT- catalase; POX- peroxidase; SOD- Superoxide dismutase.

**Fig. 1a-f.**
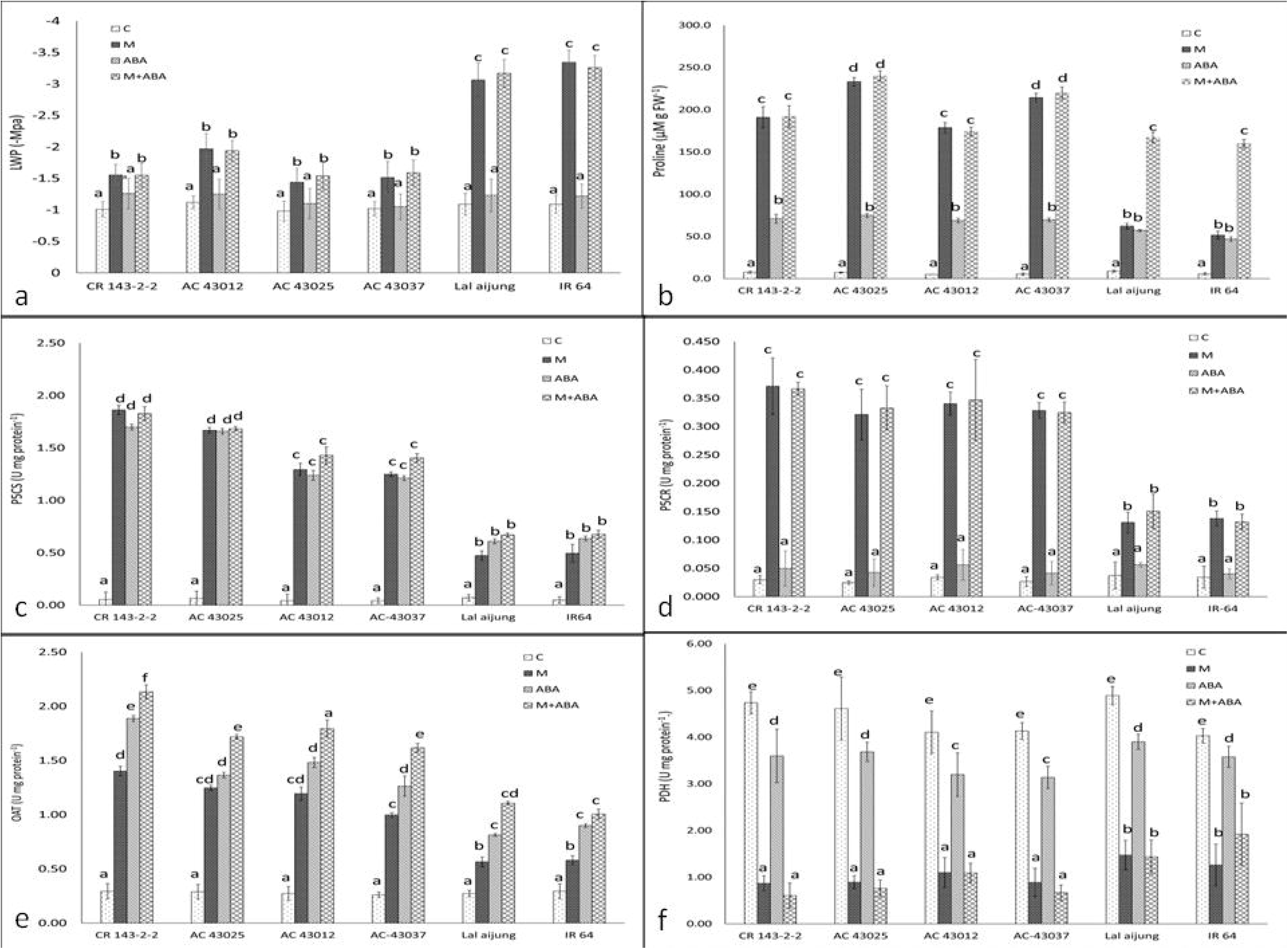
Effect of osmotic stress, ABA and combined ABA and osmotic stress on leaf water potential (LWP), proline, and proline biosynthesis enzymes (P5CS-Δ^1^-pyrroline-5-carboxylate synthase; P5CR- Δ^1^-pyrroline-5-carboxylate reductase; OAT - Ornithine-δ-aminotransferase; PDH- proline dehydrogenase) in four drought tolerant and two drought sensitive genotypes. Error bars represent SE values (n=3). Different letters indicate significant differences (*P* < 0.05) among the treatments and within the genotypes. Control (C) corresponds to normal growth condition without any treatment. Osmotic stress (M) corresponds to 2% D-mannitol mediated osmotic stress; ABA corresponds to 10 μM of ABA applied exogenously through the nutrient medium; M+ABA corresponds to 2% D-mannitol and 10 μM ABA applied simultaneously to the nutrient medium.

### Effect of osmotic stress and ABA on proline content and proline biosynthesis enzymes

Proline content increased significantly (p<0.05) under all the three treatments (M, ABA and M+ABA) compared to control. Accumulation of proline was increased up to 22-fold and 27-fold under M and M+ABA treatment respectively after six days of treatments (Fig. 1). In ABA treatment, the accumulation of proline was only 10-fold compared to control. However, proline accumulation was more in M+ABA compared to osmotic stress (M) and ABA treatment. Significant variation was observed in proline content (p<0.05) between tolerant and susceptible genotypes. Variation in proline content in susceptible genotypes Lalaijung and IR 64 was insignificant in M and ABA treatments, however, in M+ABA treatment, Lalaijung had significantly higher proline content compared to IR 64. To elucidate the differential accumulation of proline under the three treatments, the rate of activity of enzymes involved in proline biosynthesis was studied. The activity of P5CS was 20-fold more as compared to control under all the three treatments after six days of treatment. P5CS activity was significantly lower in ABA treatments of all tolerant genotypes compared to their corresponding M and M+ABA treatments. In susceptible genotypes, P5CS activity was higher than M (p<0.05) but lower than M+ABA (p<0.05). Amount of proline accumulated in susceptible genotypes was much lower than tolerant genotypes. (Fig. 1).

P5CR activity was significantly (p<0.05) higher in M and M+ABA compared to C and ABA (7 to 9-fold increase). No significant variation in P5CR activity was observed neither between C and ABA nor M and M+ABA. However, P5CR activity in tolerant genotypes were significantly higher than in susceptible genotypes. (Table 1; Fig.1).

OAT activity varied significantly in all the treatments (p<0.05). OAT activity increased about three-fold under all the three treatments compared to control after six days of treatment. OAT activity was higher in ABA compared to M but lower than M+ABA.

PDH decreased drastically in all the three treatments after six days of imposing treatments. M and M+ABA caused 60-78% reduction in PDH activity whereas ABA alone reduced only 20% of the enzyme activity depicted that PDH activity is less influenced by ABA. (Table 1; Fig. 1).

### Effect of omotic stress and ABA on anti-oxidative enzymes

Antioxidative enzyme activity was increased under all the three treatments compared to control. On an average, catalase activity was increased more than 2.5 times under all the three treatments compared to control. A significant difference in catalase activity between tolerant and susceptible genotypes was observed (Table 1). Increase in catalase activity was approximately 60% more under osmotic stress and 16-23% more under ABA and M+ABA in tolerant genotypes compared to susceptible ones. (Table 1; Fig. 2).

**Fig. 2a-c.**
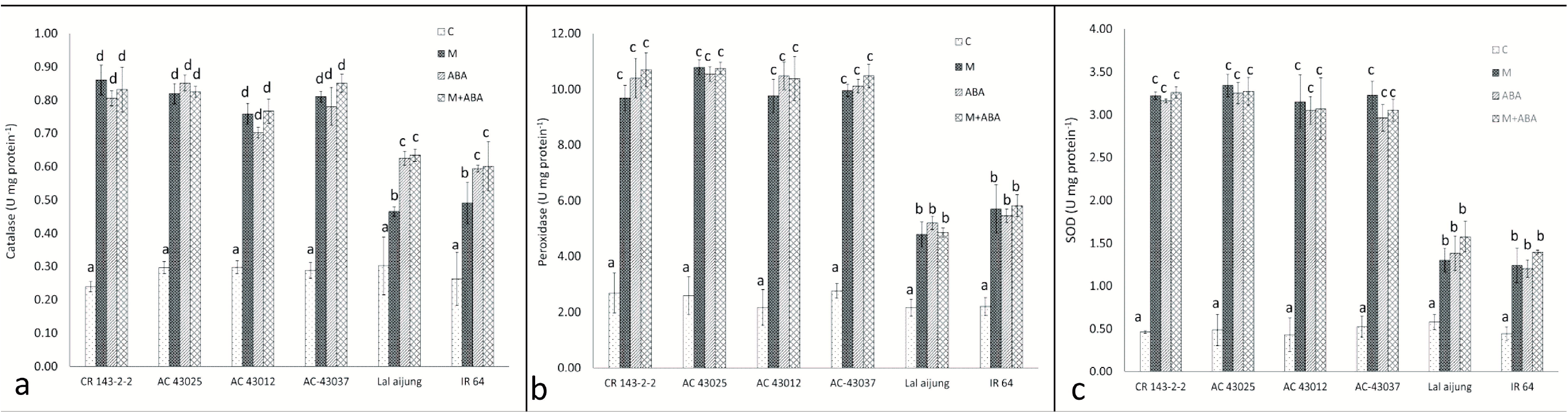
Effect of osmotic stress, ABA and combined ABA and osmotic stress on catalase (CAT), peroxidae (POX) and superoxide dismutase (SOD) in four drought tolerant and two drought sensitive genotypes. Error bars represent SE values (n=3). Different letters indicate significant differences (*P* < 0.05) among the treatments and within the genotypes. Control (C) corresponds to normal growth condition without any treatment. Osmotic stress (M) corresponds to 2% D-mannitol mediated osmotic stress; ABA corresponds to 10 μM of ABA applied exogenously through the nutrient medium; M+ABA corresponds to 2% D-mannitol and 10 μM ABA applied simultaneously to the nutrient medium.

Similarly, peroxidase activity was increased under all the three treatments and reached up to three-fold compared to control. Tolerant genotypes showed about 2-fold increase in peroxidase activity over susceptible genotypes under M, ABA and M+ABA. (Table 2 & 4; Fig 3).

**Fig. 3.**
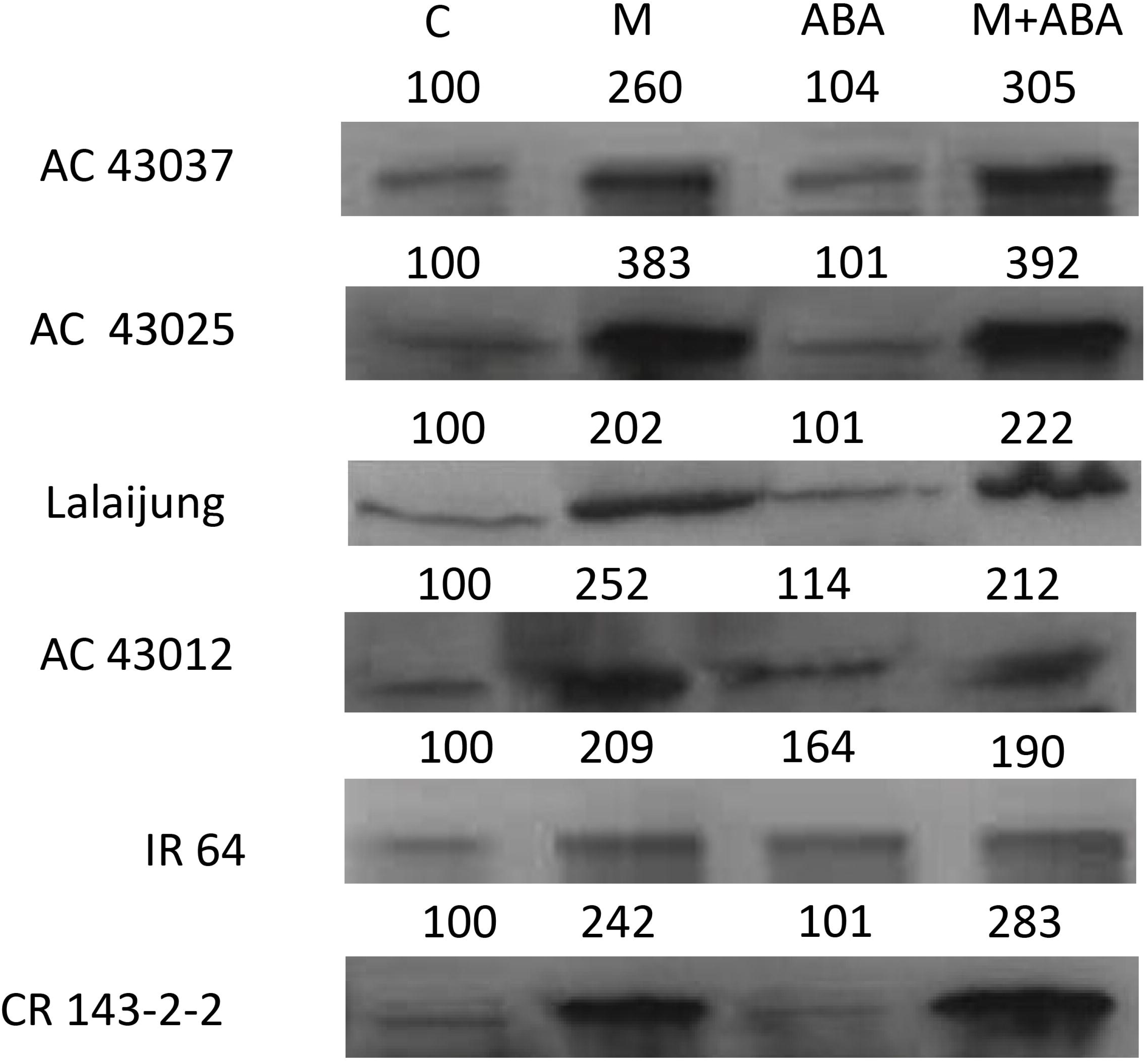
Zymograms of aldehyde oxidase (AO) activity in leaves of six rice genotypes exposed to osmotic stress, ABA and combined ABA and osmotic stress for 6 days grown in the nutrient solution. Aldehyde oxidase activity was developed using indole-3-aldehyde as substrate after native PAGE. Each lane in the gel was loaded with 400 μg of leaf proteins. Aldehyde oxidase activity is represented as the percentage after scanning and analyzing using NIH ImageJ 1.6 software assuming activity under control as 100%. Control (C) corresponds to normal growth condition without any treatment. Osmotic stress (M) corresponds to 2% D-mannitol mediated osmotic stress; ABA corresponds to 10 μM of ABA applied exogenously through the nutrient medium; M+ABA corresponds to 2% D-mannitol and 10 μM ABA applied simultaneously to the nutrient medium.

Superoxide dismutase activity was also increased under all the three treatments and amounting about 5-fold increase under M and M+ABA. Under ABA treatment, 4-fold increase in SOD activity was recorded after six days of treatment. It was also observed that under osmotic stress, tolerant genotypes showed 2.4-fold increase in SOD activity than susceptible genotypes whereas under ABA and M+ABA treatments, SOD activity increased about 1.2-fold in tolerant genotypes (Table 1; Fig. 2).

### Effect of osmotic stress and ABA on aldehyde oxidase activity

A significant increase in AO activity was observed under M and M+ABA but not under ABA treatment. Mean AO activity was increased up to 258.6% after 6 days of treatment under M whereas, under M+ABA, it was increased up to 271.3%. Under ABA treatment increase in mean AO activity was insignificant (128%) compared to M and M+ABA. It clearly indicates that AO activity is stimulated in response to osmotic stress but had no response in presence of ABA (Fig-3).

## Discussion

Earlier studies reported the active involvement of ABA during osmotic stress. The present study represents how ABA modulates different physiological and biochemical traits during osmotic stress. In many reports, ABA is known to induce stress tolerance during osmotic stress at very low concentration through alteration at the physiological as well as at molecular level but at high concentrations, it induces oxidative stress (Kuo and Ching 2004). In our study, the decrease in LWP during osmotic stress (M) has been observed that may be due to rapid transpiration and slower water absorption through roots. The effect of ABA during stress may help in maintaining LWP through closure of stomata. No such variation in LWP was observed between M and M+ABA that clearly indicate that ABA has no influence on LWP. However, a significant difference in LWP (p<0.05) was observed between tolerant and susceptible genotypes. Decrease in LWP in susceptible genotypes (Lalaijung and IR 64) was more than double to that of tolerant genotypes (AC 43025, AC 43012, AC 43037 and CR 143-2-2) both under M and M+ABA treatments that indicates rapid transpiration in susceptible genotypes resulting in loss of turgidity and hence very low water potential compared to tolerant genotypes. Maintenance of high LWP in response to moisture stress can be the result of the accumulation of compatible solutes like proline in tolerant genotypes, that decreases the cellular osmotic potential to prevent water loss (Zhang et al. 1999). In our study, an increase in proline content was less in ABA compared to osmotic stress (M). Proline accumulation might be independent of the ABA-mediated signaling pathway. According to Zhang et al. (1999), endogenous ABA accumulation is required for proline accumulation but ABA alone is not sufficient to elicit the levels of proline accumulation observed under low water potential. The reason for this differential response might be due to the response of biosynthesis enzymes involved in proline biosynthesis towards ABA. According to Savoure et al. (1997), both the expression of P5CS and P5CR were up-regulated under osmotic stress but exogenous ABA application enhanced the expression of P5CS but not P5CR. Therefore, the glutamate might be converted to glutamate-γ-semialdehyde (GSA) but unable to form proline due to reduced expression of enzyme P5CR. This supported our finding where a 24-fold increase in P5CS activity was observed in osmotic stress (M) and the combined effect of osmotic stress and ABA (M+ABA). Under ABA treatment, a 21-fold increase in P5CS activity was observed which was almost equal to the amount of P5CS activity produced under other two treatments (M and M+ABA). The activity of P5CR was insignificant from that of the control under the treatment of ABA. However, under osmotic stress (M) and combined effect of osmotic stress and ABA (M+ABA), P5CR activity was enhanced leading to an increase in proline content to more than 20-fold increase. The alternative pathway to proline production catalyzed by OAT also enhanced both in response to osmotic stress and exogenously applied ABA. Many authors suggested that *OAT* can also produce P5C and contribute to proline accumulation (Roosens et al. 2002). Funck et al, (2008) discarded the involvement of OAT in proline accumulation and proline biosynthesis occurred predominantly via the glutamate pathway in Arabidopsis. However, the function of OAT in Arabidopsis has been questioned in part because of new information showing that it is localized in the mitochondria (Funck et al. 2008). This changing role of OAT makes it of even greater importance to determine its regulation under conditions leading to proline accumulation. Sharma and Verslues (2010) stated that exogenous ABA did not induce OAT expression which was against our findings in which 3.5-fold increase in OAT activity was observed in response to ABA. Insignificant difference in OAT activity in M, ABA and M treatments gave insights on the regulation of OAT activity by both ABA dependent and ABA independent pathways. Moreover, it can be inferred that reduced activity of P5CR might have resulted in reduced proline synthesis under ABA treatment and most of the accumulated proline might have contributed by OAT activity. The difference in proline accumulation between M and M+ABA treatment might be due to the contribution of OAT in response to ABA under M+ABA treatment. According to You et al. (2012), OAT was strongly induced by ABA in rice and both ABA-dependent and AB-independent pathway contribute to the drought-induced expression of OAT. This is in accordance with our findings. Proline dehydrogenase (PDH) is the enzyme, which is associated with the oxidation of proline to glutamate thus responsible for the reduction in proline level in cells. In our study, the activity of PDH was higher in the control condition but gradually decreased under stress. According to Shevyakova et al. (2013), ABA suppressed PDH activity when treated to the roots. Dallmier and Stewart (1992) also stated that exogenous ABA declines PDH activity, which is very small, compared to level declined in response to drought stress suggesting that ABA is not the pathway linking drought stress and PDH activity. In our experiment, no response of ABA was observed on PDH activity. However, PDH activity was declined under osmotic stress and combined effect of osmotic stress and ABA. There was much difference in PDH activity between susceptible and tolerant genotypes that might be due to lack of intracellular signal or failure in the suppression of PDH activity that contributed to low levels of proline content in susceptible genotypes.

The first response to osmotic stress is the production of free radicals that induce expression of many genes like anti-oxidative enzymes that helped in diluting the effect of osmotic or oxidative stress. ABA biosynthesis enzymes also get activated in response to free radicals that induce H_2_O_2_ production and activated anti-oxidative enzymes like catalases, peroxidases and superoxide dismutase. Interrelationship between the generation of reactive oxygen species in response to PEG-induced osmotic stress and in response to ABA with activation of anti-oxidative enzymes has been established (Jiang and Zhang 2002). This is in accordance with our finding where peroxidase, catalase and superoxide dismutase activity were enhanced by in response to M, ABA and M+ABA. Rivero et al., (2007) also reported an increase in the transcript level of SOD in transgenic lines of tobacco plants under drought. According to the findings of Rivero et al., (2007), efficient scavenging of ROS protects the photosynthetic apparatus during drought stress that leads to improved water use efficiency during and after drought stress.

The plant hormone abscisic acid has long been known to be involved in the response of plants to various environmental stresses, particularly drought (Zhu 2011). It mediates coordination between growth and development in response to the environment. Under non-stressful conditions, low ABA level is maintained in the plant cell which is required for normal plant growth (Finkelstein and Rock 2002). In this investigation, the effect of osmotic stress, ABA and the combined effect of osmotic stress and ABA (M+ABA) on the activity of aldehyde oxidase has been studied. The activity of AO was enhanced in response to osmotic stress and under the combined effect of osmotic stress and ABA. According to Xiong and Zhu (2003), the degradation of ABA is suppressed by osmotic stress and activated by ABA and stress relief which is in accordance with our findings. No significant change in AO activity was observed under ABA treatment. Qin and Zeevaart (2002) have demonstrated that overproduction of ABA correlated with the over-accumulation of phaseic acid which leads to the notion that ABA might restrict its own accumulation by activating its degradation even under non-stressful conditions.

## Conclusion

Abscisic acid is a phytohormone that positively regulates drought tolerance at low concentration but induces senescence at high concentration. Though activation of many drought responsive enzymes is ABA-dependent, some are regulated independently of ABA. The present study revealed how ABA modulates different biochemical traits that are involved in the drought tolerance mechanism which signifies the role of ABA in drought tolerance mechanism and provided information for the development of an effective drought management strategy. Though P5CS and OAT are activated through ABA, P5CR and PDH remain unresponsive which indicates proline can be synthesized independently of ABA from glutamate or can be synthesized from ornithine through an ABA-dependent pathway. Anti-oxidative enzymes produced through ABA-dependent pathway that helps in ameliorating the effects of drought stress. ABA also regulates its own endogenous level by down-regulating its own activity to maintain its optimal level inside the cell. The information provided here will be useful for carrying out an extensive analysis of the role of ABA in drought tolerance.

## Acknowledgment

Authors are thankful to the NICRA (National Innovation on Climate Resilient Agriculture) project for providing financial support to conduct this experiment.

## Abbreviations

P5CS: Δ^1^-pyrroline-5-carboxylate synthase
P5CR: Δ^1^-pyrroline-5-carboxylate reductase
OAT: Ornithine-δ-aminotransferase
PDH: proline dehydrogenase
M: Mannitol
ABA: Abscisic acid
LWP: Leaf water potential
AO: Aldehyde oxidase
MTT: 3[4,5-dimethylthiazol-2-yl]-2,5-diphenyltetrazolium-bromide
GSH: reduced glutathione
DTT: dithiothreitol
PAGE: polyacrylamide gel electrophoresis

## References

Bates LS, Waldren RP, Teare ID (1973) Rapid determination of free proline for water stress studies. Plant Soil 207:205–207

Dallmier KA, Stewart CR (1992) Effect of exogenous abscisic acid on proline dehydrogenase activity in maize (Zea mays L.). Plant Physiol 99:762–764. doi: 10.1104/pp.99.2.762

Dash GK, Barik M, Debata AK, et al (2017) Identification of most important rice root morphological markers in response to contrasting moisture regimes under vegetative stage drought. Acta Physiol Plant 39:. doi: 10.1007/s11738-016-2297-1

Dash GK, Swain P (2015) Differential response of rice genotypes to mild and severe osmotic stress during seedling stage. Oryza 52:307–312

Farooq M, Kobayashi N, Ito O, et al (2010) Broader leaves result in better performance of indica rice under drought stress. J Plant Physiol 167:1066–1075. doi: 10.1016/j.jplph.2010.03.003

Finkelstein RR, Rock CD (2002) Abscisic Acid Biosynthesis and Response. Arab B 1:e0058. doi: 10.1199/tab.0058

Funck D, Stadelhofer B, Koch W (2008) Ornithine-δ-aminotransferase is essential for Arginine Catabolism.pdf. BMC Plant Biol 8:1–14

Giannopolitis CN, Ries SK (1977) Superoxide dismutases: I. Occurrence in higher plants. Plant Physiol 59:309–30914. doi: 10.1104/pp.59.2.309

H.D. Barrs and P.E. Weatherley (1962) A Re-Examination of the Relative Turgidity Technique for Estimating Water Deficits in Leaves. Aust J Biol Sci 15:413–428. doi: https://doi.org/10.1071/BI9620413

Jiang M, Zhang J (2001) Effect of abscisic acid on active oxygen species, antioxidative defence system and oxidative damage in leaves of maize seedlings. Plant Cell Physiol 42:1265–1273. doi: 10.1093/pcp/pce162

Jiang M, Zhang J (2002) Water stress-induced abscisic acid accumulation triggers the increased generation of reactive oxygen species and up-regulates the activities of antioxidant enzymes in maize leaves. J Exp Bot 53:2401–2410. doi: 10.1093/jxb/erf090

Jiang W, Bai J, Yang X, et al (2012) Exogenous application of abscisic acid, putrescine, or 2,4-epibrassinolide at appropriate concentrations effectively alleviate damage to tomato seedlings from suboptimal temperature stress. Horttechnology 22:137–144

Kholová J, Hash CT, Kumar PL, et al (2010) Terminal drought-tolerant pearl millet [Pennisetum glaucum (L.) R. Br.] have high leaf ABA and limit transpiration at high vapour pressure deficit. J Exp Bot 61:1431–1440. doi: 10.1093/jxb/erq013

Kuo TH, Ching HK (2004) Hydrogen peroxide is necessary for abscisic acid-induced senescence of rice leaves. J Plant Physiol 161:1347–1357. doi: 10.1016/j.jplph.2004.05.011

Laemmli UK (1970) Cleavage of Structural Proteins during the Assembly of the Head of Bacteriophage T4. Nature 227:680–685

Lian HL, Yu X, Lane D, et al (2006) Upland rice and lowland rice exhibited different PIP expression under water deficit and ABA treatment. Cell Res 16:651–660. doi: 10.1038/sj.cr.7310068

Lowry OH, Rosebrough NJ, Farr AL, Randall RJ (1951) Protein Measurement with the Folin Phenol Reagent. J Biol Chem 193:265–275

Lutts S, Majerus V, Kinet J (1999) NaCl effects on proline metabolism in rice.pdf. Physiol Plant 105:450–458

Madan S, Nainawatee HS, Jain RK, Chowdhury JB (1995) Proline and proline metabolising enzymes in in-Vitro selected NaCl-Tolerant Brassica juncea L. Under salt stress. Ann. Bot. 76:51–57

Pereira GJG, Molina SMG, Lea PJ, Azevedo RA (2002) Activity of antioxidant enzymes in response to cadmium in Crotalaria juncea. Plant Soil 239:123–132. doi: 10.3923/jbs.2009.44.50

Pütter J (1974) Peroxidases. In: Methods of Enzymatic Analysis

Qin X, Zeevaart JAD (2002) Overexpression of a 9-cis-Epoxycarotenoid Dioxygenase Gene in. J Biol Chem 128:544–551. doi: 10.1104/pp.010663.The

Rivero RM, Kojima M, Gepstein A, et al (2007) Delayed leaf senescence induces extreme drought tolerance in a flowering plant. Proc Natl Acad Sci U S A 104:19631–19636. doi: 10.1073/pnas.0709453104

Roosens NH, Bitar F Al, Loenders K, et al (2002) Overexpression of ornithine-δ-aminotransferase increases proline biosynthesis and confers osmotolerance in transgenic plants. Mol Breed 9:73–80. doi: 10.1023/A:1026791932238

Rothe GM (1974) Aldehyde oxidase isoenzymes (E.G. 1. 2. 3. 1) in potato tubers (Solanum tuberosum). Plant Cell Physiol 15:493–499

Sansberro PA, Mroginski LA, Bottini R (2004) Foliar sprays with ABA promote growth of Ilex paraguariensis by alleviating diurnal water stress. Plant Growth Regul 42:105–111. doi: 10.1023/B:GROW.0000017476.12491.02

Sarfaraz-Ardakani M-R, Khavari-Nejad R-A, Moradi F, Najafi F (2014) Abscisic acid and cytokinin-induced carbohydrate and antioxidant levels regulation in drought-resistant and -susceptible wheat cultivar during grain filling under field conditions. Int J Biosci 5:11–24. doi: 10.12692/ijb/5.8.11-24

Savoure A, Hua XJ, Bertauche N, et al (1997) Abscisic acid-independent and abscisic acid-dependent regulation of proline b.pdf. Mol Gen Genet 254:104–109

Sharma S, Verslues PE (2010) Mechanisms independent of abscisic acid (ABA) or proline feedback have a predominant role in transcriptional regulation of proline metabolism during low water potential and stress recovery. Plant, Cell Environ 33:1838–1851. doi: 10.1111/j.1365-3040.2010.02188.x

Shevyakova NI, Musatenko LI, Stetsenko LA, et al (2013) Effect of ABA on the contents of proline, polyamines, and cytokinins in the common ice plants under salt stress. Russ J Plant Physiol 60:741–748. doi: 10.1134/S1021443713060125

Sibounheuang V, Basnayake J, Fukai S (2006) Genotypic consistency in the expression of leaf water potential in rice (Oryza sativa L.). F Crop Res 97:142–154. doi: 10.1016/j.fcr.2005.09.006

Stines AP, Naylor DJ, Høj PB, Van Heeswijck R (1999) Proline accumulation in developing grapevine fruit occurs independently of changes in the levels of δ1-pyrroline-5-carboxylate synthetase mRNA or protein. Plant Physiol 120:923–931

Turner N (1982) The role of shoot characteristics in drought resistance of crop plants. Drought Resist Crop with Emphas rice 115–134

Vogel RH, Kopac MJ (1960) Some properties of ornithine δ-transaminase from Neurospora. BBA - Biochim Biophys Acta. doi: 10.1016/0006-3002(60)90517-5

Xiong L, Zhu J-K (2003) Update on abscisic acid biosynthesis. Plant Physiol 133:29–36. doi: 10.1104/pp.103.025395.mutant

Xoconostle-Cazares B, Ramirez-Ortega FA, Flores-Elenes L, Ruiz-Medrano R (2010) Drought Tolerance in Crop Plants Review Molecular.Pdf. Am J Plant Physiol 5:241–256

Yoshida S, Forno, Douglas A.Cock JH, Gomez K a (1976) Laboratory Manual for Physiological Studies of Rice, 3rd edn. The International Rice Research Institute, Los Banas, Philippines

You J, Hu H, Xiong L (2012) An ornithine δ-aminotransferase gene OsOAT confers drought and oxidative stress tolerance in rice. Plant Sci 197:59–69. doi: 10.1016/j.plantsci.2012.09.002

Zdunek E, Lips SH (2001) Transport and accumulation rates of abscisic acid and aldehyde oxidase activity in Pisum sativum L. in response to suboptimal growth conditions. J Exp Bot 52:1269–1276. doi: 10.1093/jxb/52.359.1269

Zhang J, Jia W, Yang J, Ismail AM (2006) Role of ABA in integrating plant responses to drought and salt stresses. F Crop Res 97:111–119. doi: 10.1016/j.fcr.2005.08.018

Zhang J, Nguyen HT, Blum A (1999) Genetic analysis of osmotic adjustment in crop plants. J Exp Bot 50:291–302. doi: 10.1093/jxb/50.332.291

Zhu J (2011) Salt and Drought Stress Signal Transduction in Plants. Annu Rev Plant Biol 53:247–273. doi: 10.1146/annurev.arplant.53.091401.143329.Salt

